# A Validated LC–MS/MS Method for Quantifying Phenolic Acids, Lignans, and Enterolignans from Human Fecal Samples

**DOI:** 10.1101/2025.09.01.673557

**Authors:** Christopher A. Dicksion, Darrian N. Chao, Jonathan D. Rickmeyer, Elizabeth N. Bess

## Abstract

The human gut microbiome is home to numerous small molecules that impact health. Three prominent classes of molecules in this environment include phenolic acids, lignans, and enterolignans, which have been linked to anti-inflammatory and anti-oxidant effects as well as protection from cancer, cardiovascular disease, and neurodegeneration. The abundance of these molecules in the gut microbiome as well as their biological significance motivated the development of the LC–MS/MS method reported herein, which provides a simple, robust, and high-throughput approach to simultaneously quantify a 17-membered panel of phenolic acids, lignans, and enterolignans in human fecal microbiome samples. This method employs liquid-liquid extraction, which allows for cost-effective sample preparation by using common laboratory materials. Additionally, fecal samples are lyophilized prior to extraction to mitigate confounding effects introduced by variability in water content that exists across individuals’ samples. Inclusion of 0.1% acetic acid in the chromatographic solvents optimized peak shape and signal intensity. Plastic microcentrifuge tubes imparted substantial interferences for some analytes, which was resolved through use of glass vials. Using this method, 60% of human fecal samples from 10 individuals showed that phenolic acids bearing a catechol motif were in significantly greater abundance than were guaiacols. As specific gut bacteria can transform guaiacols into redox-active catechols, determination of guaiacol:catechol ratios in fecal samples may afford a biomarker of gut microbiota anti-oxidant potential.

**For the Table of Contents only:** 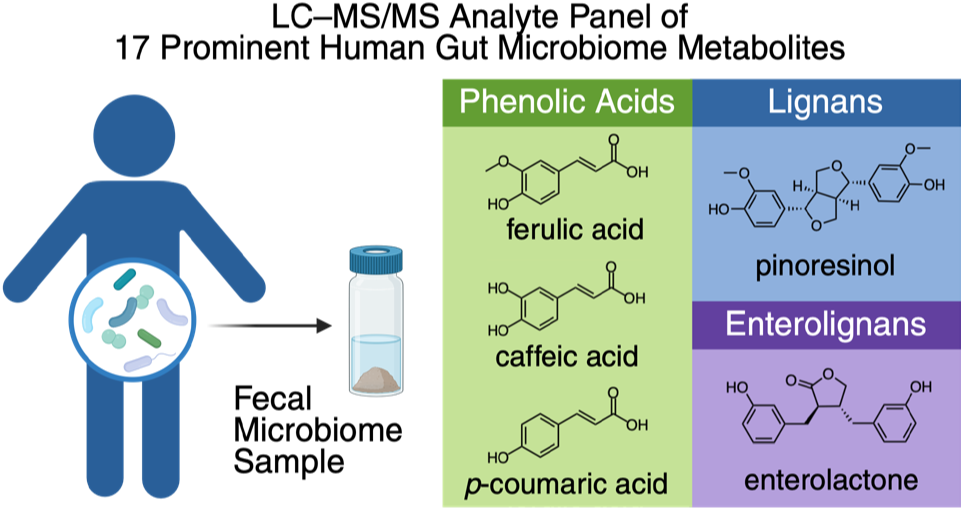

## INTRODUCTION

Metabolites produced by bacteria in the human gut are emerging as critical modulators of human health.^1,2^ As such, the need for analytical methods to identify and quantify metabolites produced by gut bacteria is critical for studies that seek to link metabolomic profiles to the development, progression, and treatment of disease.^3,4,5^ Equally important is the development of methods for the quantitation of health-promoting gut microbial metabolites, such as the diverse class of polyphenols, including phenolic acids, lignans, and enterolignans.^6^ These compound classes contain molecules that exert antioxidant,^7^ anti-inflammatory,^8^ antihypertensive,^9^ and anticancer effects,^10^ highlighting their potential as therapeutic agents against a variety of diseases. To facilitate studies that seek to identify new links between gut bacterial metabolism and human health, we have validated an LC–MS/MS method for the fecal-sample quantification of a 17-membered panel of molecules that are derived from plant-based diets: 10 phenolic acids, 5 lignans, and 2 enterolignans. Phenolic acids and lignans are biosynthesized in plants from the same molecular precursors,^11^ and enterolignans are synthesized from lignans in human gut microbiotas.^12^ The linked origins of each molecule in this panel as well as their prevalence^13^ and abundance^14^ in human gut microbiotas motivated us to develop this analytical method to simultaneously quantify each analyte in this collection.

The largest class of molecules measured by our method is phenolic acids. These plant secondary-metabolites (**Figure 1A**) are prevalent in fruits, vegetables, and beverages like coffee and tea.^15^ Average dietary intake of phenolic acids among U.S. adults is estimated to be 1005.6 mg per day.^16^ Phenolic acids not only contribute to the flavor, aroma, and nutritional profiles of foods and beverages,^17^ but they also have properties that may fight diseases ranging from cancer to neurodegeneration. For example, Balsamo et al. showed that one of the most diet-abundant phenolic acids, caffeic acid,^14^ significantly limits a Parkinsonian neurodegenerative pathway (i.e., formation of ɑ-synuclein aggregates) in murine enteroendocrine cells of the intestine; caffeic acid exerts this effect by suppressing the oxidation of dopamine, which mediates this pathogenic process.^18^ Zhang et al. showed that another prominent diet-derived phenolic acid, ferulic acid,^19^ not only inhibited metastasis of MDA-MB-231 breast cancer cells in vitro, but it also decreased tumor volume and increased apoptosis of tumor cells that were xenografted into mice.^20^ Another member of this compound class—3-(3-hydroxyphenyl)propanoic acid (3,3-HPPA), which is particularly abundant in legumes and whole grains—decreased arterial blood pressure in spontaneously hypertensive rats, suggesting that its vasodilatory effects promote cardiovascular health.^9^

**Figure 1:**
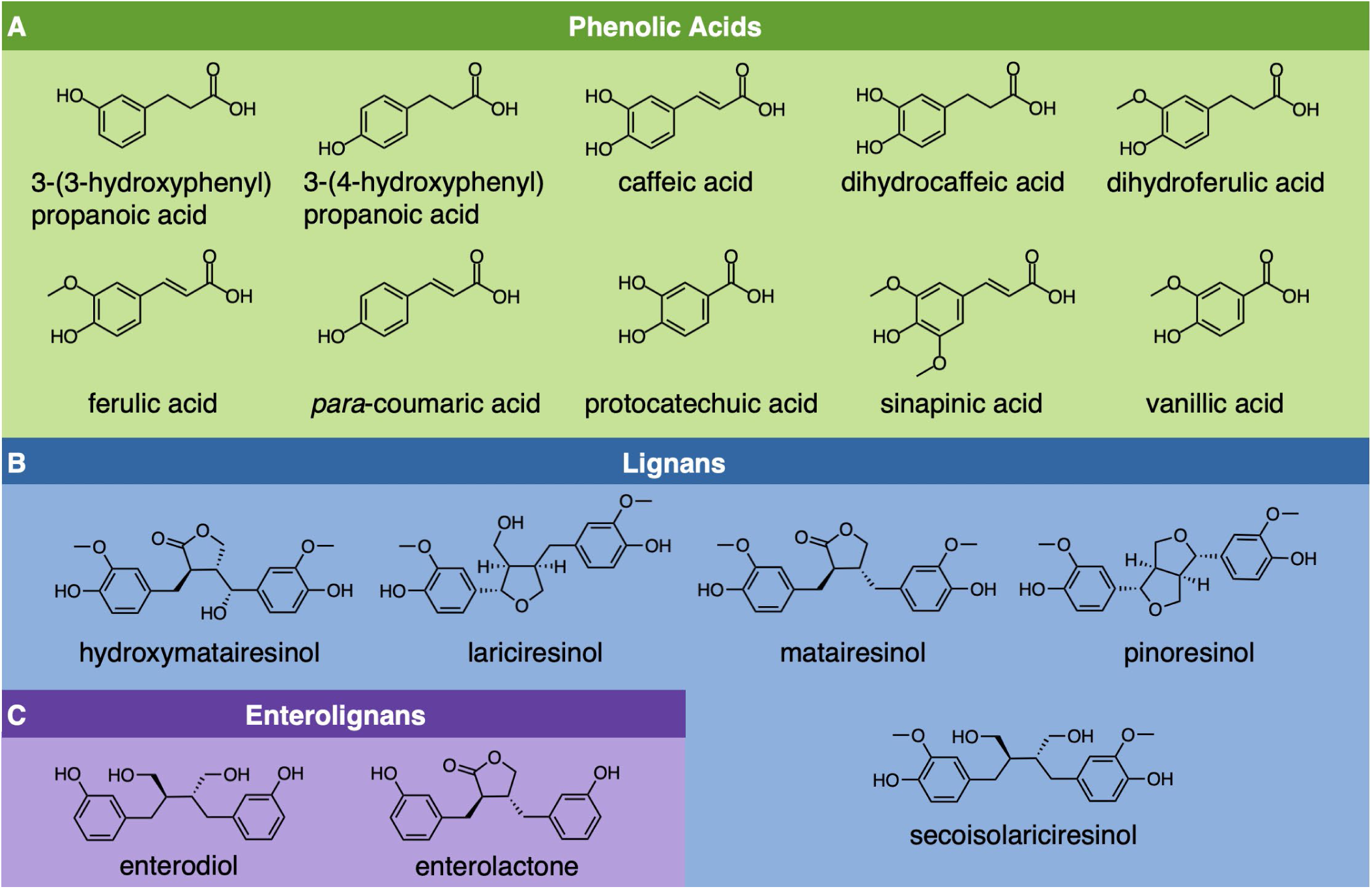
Analyte Panel. Collection of the ten phenolic acids, five lignans, and two enterolignans quantified using the LC–MS/MS method reported herein.

The next class of molecules measured using the method reported here is lignans (**Figure 1B**). These dimeric phenylpropanoid metabolites are found in fiber-rich foods, especially seeds and nuts,^21^ with total dietary consumption averaging around 1 mg per day.^22^ Lignans have been shown to exert anticancer effects by inhibiting tumor growth and metastasis in animal models^23^ as well as by inducing apoptosis in breast cancer, colorectal cancer, and prostate cancer cell lines.^24^ Rats fed a diet of one such lignan, lariciresinol (LARI), showed decreased angiogenesis and increased apoptosis of tumor cells in two different rat models of breast cancer, one of dimethylbenzyl[a]anthracene-induced mammary tumors and the other of xenograft MCF-7 breast cancer cells.^25^ Lignans may also improve cardiovascular health. A prospective study of over 200,000 U.S. adults found that increased lignan intake, especially from dietary fiber, was associated with a lower risk of coronary heart disease.^26^

In addition to diet-derived lignans exerting health-promoting effects, they are also metabolized by a consortium of human gut bacteria into enterolignans, namely enterodiol (END) and enterolactone (ENL) (**Figure 1C**).^27^ Enterolignans, which are quantified via the method reported here, exhibit their own distinct anticancer effects.^10^ When gnotobiotic rat models of breast cancer were colonized with a lignan-metabolizing bacterial consortium and fed a lignan-rich diet, fecal concentrations of ENL increased in tandem with tumor-cell apoptosis and overall tumor burden decreased as compared with uncolonized gnotobiotic rats.^28^ Aligned with these findings, studies in the MCF-7 breast cancer cell line show that ENL is an aromatase inhibitor that limits proliferation of cancer cells by interrupting the formation of estradiol.^29^ Taken together, these findings suggest that gut microbiome profiles encoding for specific biosynthetic pathways may impact breast cancer risk and improve health outcomes for people living with breast cancer. Indeed, a recent meta-analysis of three observational studies encompassing 3,864 participants concluded that increased serum ENL concentration is correlated with decreased mortality from breast cancer.^30^

Due to the health-promoting effects of phenolic acids, lignans, and enterolignans, we selected a panel of 17 analytes that are linked by their biosynthetic origins (**Figure 1**) and sought to quantify each within a single LC–MS/MS method. While others have reported several methods that quantify subsets of these compounds, only one other method simultaneously captures them all; however the utility of this method is hampered by its 14-minute run-time.^31^ An early method, published in 1966 by Dalgliesh and coworkers, used GC–MS to quantify a wide variety of phenolic acids in urine, paving the way for metabolomics studies that investigated the impact of phenolic acids on human health.^32^ It was not until 2003 when Gonthier et al. published an LC–MS/MS method for the quantitation of numerous hydroxycinnamic acids found in urine.^33^ In 2011, Sánchez-Patán et al. published an LC–MS/MS method to quantify an extensive panel of 47 phenolic acid metabolites in fecal samples; however, the total run-time of 18 minutes per injection precluded its application to metabolomics studies on large sample sets.^34^

Methods for the quantitation of lignans and enterolignans date back to 1982, when Fotsis et al. developed a GC–MS method to quantify END and ENL in urine samples.^35^ Adlercreutz and colleagues applied the Fotsis method to urine samples of postmenopausal women, finding that women with breast cancer excreted lower urinary concentrations of ENL than did healthy women, thereby setting the stage for method development to identify and quantify lignans and enterolignans in human samples.^36^ Heinonen and coworkers expanded this method in 2001 to quantify 11 lignan metabolites in fecal samples; however, the extensive sample preparation steps, including the requisite derivatization step with a silylating agent for GC–MS compatibility, spurred the development of LC–MS/MS approaches to quantify lignans from biological matrices.^37^ In 2006, Smeds et al. pioneered an LC–MS/MS method to quantify 5 lignans and 3 enterolignans from human serum.^38^ In 2010, Prasain and colleagues published a fast LC–MS/MS method—2-minute run-time per injection—for the quantification of END and ENL from plasma, but the method does not include any coverage of the precursor metabolites to these enterolignans.^39^ More recently, Nørskov and Knudsen published an LC–MS/MS method for the quantitation of 10 lignans in pig fecal samples, including both enterolignans, with a modest run-time per injection of 7.8 min.^40^

The targeted LC–MS/MS method presented herein provides a 3-pronged contribution to the science of gut microbiome metabolomics, and it has been validated to conform to established FDA guidelines.^41^ First, whereas other targeted mass spectrometric methods can quantify either phenolic acids, lignans, or enterolignans, the current method can account for analytes from all three classes of compounds in a single injection. The method developed by González-Domínguez et al. also quantified molecules from these 3 classes in a single injection, but the sample preparation was optimized for plasma, the run-time per injection was 14 minutes, and the lower limit of quantitation for the lignans and enterolignans was 16-fold and 26-fold greater, respectively, than what is achieved by the new method reported here.^31^ Second, our method normalizes analyte concentrations to the mass of dry fecal samples; due to differences in the water content of fecal samples from person to person, normalizing the analyte content to the dry mass of the fecal sample provides a reliable and reproducible way to standardize metabolite abundance comparisons between individuals, across studies, and over time.^42,43^ Lastly, this method contributes enhanced analytical throughput; whereas most existing methods for these analytes have total run-times between 8 and 28 minutes, the run-time per sample for the method reported here is only 5 minutes.

## MATERIALS AND METHODS

### Chemicals

Caffeic acid, *para*-coumaric acid, and protocatechuic acid were purchased from Tokyo Chemical Industry, Co., in 98% purity. Ferulic acid and 3-(4-hydroxyphenyl)propanoic acid were purchased from Sigma Aldrich, in 99% and 98% purity, respectively. Dihydrocaffeic acid and 3-(3-hydroxyphenyl)propanoic acid were purchased from Alfa Aesar in 98% purity. Dihydroferulic acid was purchased from Matrix Scientific in 98% purity. Sinapinic acid was purchased from AK Scientific in 95% purity. Vanillic acid was purchased from Sigma Aldrich in 97% purity. (–)-Matairesinol was purchased from Santa Cruz Biotechnology in 95% purity. (–)-Enterodiol was purchased from PhytoLab in 95% purity. (–)-Enterolactone, (–)-hydroxymatairesinol, (+)-lariciresinol, (+)-pinoresinol, and (–)-secoisolariciresinol were purchased from Sigma Aldrich in 95% purity. The internal standards (IS) ^13^C_3_-caffeic acid, d_6_-rac-secoisolariciresinol, and ^13^C_3_-rac-enterolactone were purchased from Toronto Research Chemicals in 95% purity. HPLC grade methyl *tert*-butyl ether (MtBE) as well as LC–MS grade acetonitrile and acetic acid were purchased from Fisher Scientific. Double deionized water (DDI H_2_O) was obtained from an Elga PURELAB flex water purifier at greater than 18.2 MΩ⋅cm.

### Fecal samples

Fecal samples were obtained from healthy human volunteers who consented to participate in the study, which was approved by the University of California, Irvine Institutional Review Board. Samples were collected using a GutAlive anaerobic microbiome kit (Microviable, Gijón, Spain). Upon receipt, the samples were brought into an anaerobic chamber (Coy Laboratory Products, Grass Lake, MI) with a gaseous atmosphere of 20% CO_2_, 2%–5% H_2_, and the balance N_2_. Aliquots of 0.5–2.0 g of homogenized fecal material were added to pre-weighed 15 mL conical tubes. The samples were stored at -80 °C until use. For lyophilization, 0.5–2 g of homogenized fecal material was added to pre-weighed 15 mL conical tubes in an anaerobic chamber. The samples were frozen at -80 °C for 24 hr and then lyophilized for 5 days to ensure dryness. Lyophilized samples were stored at -80 °C until use.

### Calibration and Quality Control Sample Preparation

Commercially available standards for each analyte in the 17-membered panel were weighed and dissolved in acetonitrile to make 17 individual 10 mM stock solutions. Aliquots of the individual stock solutions were stored at -80 °C until use. These individual stocks were combined to make 1 mL of a 100 µM mixture of each standard in DDI H_2_O containing the internal standards (IS) 50 nM ^13^C_3_-caffeic acid (50 µL of a 10 mM stock in acetonitrile into 99.91 mL DDI H_2_O), 20 nM each of d_6_-rac-secoisolariciresinol (20 µL of a 10 mM stock in acetonitrile into 99.91 mL DDI H_2_O) and ^13^C_3_-rac-enterolactone (20 µL of a 10 mM stock in acetonitrile into 99.91 mL DDI H_2_O). Aliquots of the 100 µM stock mixture were stored at -80 °C. Upon use, an aliquot of the 100 µM stock mixture was thawed at room temperature for 5 minutes before being diluted to 1 µM with DDI H_2_O containing IS. The calibration samples were prepared as two-fold serial dilutions from 500 nM to 0.24 nM in DDI H_2_O containing IS and were used immediately.

The quality control samples were prepared in lyophilized fecal material, using standard addition. Between 1 and 5 mg of lyophilized fecal material was weighed into a 2 mL glass HPLC vial and resuspended in 1.8 mL of DDI H_2_O. The samples were heated in a water bath at 100 °C for 15 minutes to inactivate bacteria and precipitate proteins. After chilling on ice for 15 minutes, the samples were centrifuged at 2000 rpm for 15 min to pellet insoluble debris. A 300 µL aliquot of the supernatant was transferred into each of five 2 mL glass HPLC vials. Into this supernatant, a 300 µL aliquot of each analyte standard from our 17-membered panel was added at 10 nM, 40 nM, 100 nM, or 400 nM in DDI H_2_O containing 50 nM ^13^C_3_-caffeic acid as well as 20 nM each of d_6_-rac-secoisolariciresinol and ^13^C_3_-rac-enterolactone as IS. To detect any signal endogenous to the fecal material, a 300 µL aliquot of DDI H_2_O containing IS alone was spiked into an additional 300 µL aliquot of fecal supernatant.

Next, liquid-liquid extraction was performed by adding 1.2 mL MtBE + 0.1% acetic acid to each sample. The samples were mixed at 1000 rpm for 5 minutes by vortexing (**Figure S1A**), followed by centrifugation at 2000 rpm for 5 minutes (**Figure S1B**). A 900 µL aliquot of each supernatant was transferred to its own glass culture tube. Next, the samples were placed in a 40 °C water bath and dried under a stream of N_2_ for 15 minutes with a Biotage Turbovap LV (Biotage, Uppsala, Sweden). The dried residues were resuspended in 450 µL of 20% aqueous acetonitrile, capped with a butyl rubber stopper, and mixed for 15 minutes at 500 rpm (**Figure S1C**). After preparation, samples were immediately subjected to analysis via LC–MS/MS.

### Method Validation

This method was developed to conform to established FDA guidelines, so each analyte was assessed according to sensitivity, selectivity, stability, accuracy and precision, recovery, and matrix effects.^41^

#### Sensitivity

The sensitivity of the method was established by first recording the signal intensity of 12 sequential blank injections, consisting of 20% acetonitrile + 0.1% acetic acid, to establish the background signal. Following this, a 2-fold dilution series from 500 nM to 0.244 nM of a mixture of the 17 analytes was prepared in DDI H_2_O, and the lower limit of quantitation (LLOQ) was determined to be the lowest concentration in the dilution series whose absolute signal was greater than 10 times the average signal intensity produced in the replicate blank injections.

#### Selectivity

Selectivity was assessed by first injecting a mixture of all the analytes at 1.0 µM in 20% aqueous acetonitrile + 0.1% acetic acid to establish the retention time of each analyte. Diluted fecal samples were prepared according to the quality control sample procedure (vide supra), injected into the mass spectrometer, and the presence of interfering or overlapping peaks was assessed for each analyte. Cross-signal contributions were assessed by preparing standards individually and as a mixture of all standards in 20% aqueous acetonitrile + 0.1% acetic acid at 1.0 µM.

#### Matrix Effects

Matrix effects were assessed according to the method established by Matuszewski et al.^44^ Samples were prepared in triplicate according to the quality control procedure described above, with the modification that the mixture of analyte standards at 0 nM (termed Endo), 10 nM, 40 nM, 100 nM, and 400 nM (collectively termed Spike) were spiked into the matrix samples after liquid-liquid extraction. Neat mixtures in 20% aqueous acetonitrile of the analyte standards at the same concentrations (termed Neat) were used for comparison. Samples were immediately subjected to analysis via LC–MS/MS. Results were calculated via standard addition for each replicate at each concentration as the percent difference of the spiked samples to the neat samples according to Equation 1, with negative values signifying ion suppression and positive values signifying ion enhancement. Results are reported as the mean ± S.E.M.

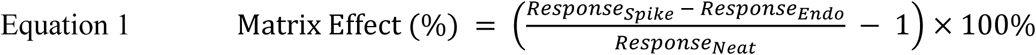

Interferences from leachate of polypropylene microcentrifuge tubes (MCT’s) were assessed by adding 600 µL of DDI H_2_O to MCT’s and extracting with MtBE + 0.1% acetic acid; sample conditions were ± 0.1% formic acid and ± heat. Two brands of 1.5 mL MCT’s and 1 brand of 2.0 mL glass HPLC vials were tested. Each condition was prepared in triplicate. The heat-treated samples were placed in a water bath at 95 °C for 15 minutes, while non-heat-treated samples were left at room temperature for an equivalent amount of time. All samples were then chilled on ice for 15 minutes, followed by liquid-liquid extraction using 1.2 mL MtBE + 0.1% acetic acid. After mixing at 1000 rpm for 5 minutes followed by centrifugation at 2000 rpm for 15 minutes, 900 µL of supernatant was transferred to a glass culture tube. The supernatants were placed in a 40 °C water bath and dried under a stream of N_2_ for 15 minutes with a Biotage Turbovap LV. The residues were resuspended in 450 µL of 20% aqueous acetonitrile, capped with a butyl rubber stopper, and mixed at 500 rpm for 15 minutes. Samples were immediately subjected to analysis via LC–MS/MS.

#### Accuracy, Precision, and Stability

Accuracy, precision, and stability measurements were made from samples prepared in the same fashion as the quality control samples (vide supra). Replicate injections (n = 6) were performed on a given sample on the same day (intraday), and independent samples were compared across multiple days (n = 6, interday). For stability studies, samples were either left on the benchtop for 12–72 hours before being analyzed, or they were frozen for 12–72 hours at -80 °C before being thawed to room temperature for 15 minutes and then analyzed. Accuracy is reported as relative error, and precision is reported as relative standard deviation.

#### Recovery

Recovery was assessed by comparing (1) the response of each analyte when a mixture of the 17 molecules in the panel was spiked into fecal matrix before liquid-liquid extraction (termed pre-extraction samples) to (2) the response of each analyte when the same mixture of 17 molecules was spiked into fecal matrix after liquid-liquid extraction (termed post-extraction samples). The samples for recovery experiments were prepared in triplicate according to the procedure described for preparing quality-control samples (vide supra). Samples were grouped as pre- and post-extraction pairs, where both used the same stock of standards and the same fecal matrix. Between 1 and 5 mg of lyophilized fecal material was weighed into a 2 mL glass HPLC vial and resuspended in 1.8 mL of DDI H_2_O. The samples were heated in a water bath at 100 °C for 15 min to kill bacteria and precipitate proteins. After chilling on ice for 15 min, the samples were centrifuged at 2000 rpm for 15 min to pellet insoluble debris. Supernatants from the paired pre-extraction and post-extraction fecal samples were combined in a glass culture tube to create a uniform endogenous mixture. A 300 µL aliquot of this mixture was transferred into each of ten 2 mL glass HPLC vials. For the pre-extraction samples, 300 µL aliquots of mixtures of analyte standards at either 0 nM (termed Endo), 10 nM, 40 nM, 100 nM, or 400 nM (termed Spike) in DDI H_2_O containing IS were spiked into the endogenous mixture. For the post-extraction samples, 300 µL of DDI H_2_O was added to the endogenous mixture. For liquid-liquid extraction, 1.2 mL MtBE + 0.1% acetic acid was added to each sample. The samples were mixed at 1000 rpm for 5 minutes, followed by centrifugation at 2000 rpm for 5 minutes. A 900 µL aliquot of each supernatant was transferred to its own glass culture tube. The samples were placed in a 40 °C water bath and dried under a stream of N_2_ for 15 minutes with a Biotage Turbovap LV. The dried residues for the pre-extraction samples were resuspended with 450 µL of 20% aqueous acetonitrile, while the post-extraction samples were resuspended with 225 µL of 40% aqueous acetonitrile and 225 µL of mixtures of analyte standards at either 0 nM (termed Endo), 10 nM, 40 nM, 100 nM, or 400 nM (termed Spike) in DDI H_2_O containing IS. All samples were capped with a butyl rubber stopper and mixed for 15 minutes at 500 rpm. Samples were subjected to analysis via LC–MS/MS immediately after preparation. Recovery was calculated via standard addition as the percentage of the measured concentration of the pre-extraction spiked samples to the concentration of the post-extraction spiked samples, according to Equation 2. The results are reported as the mean ± S.E.M.

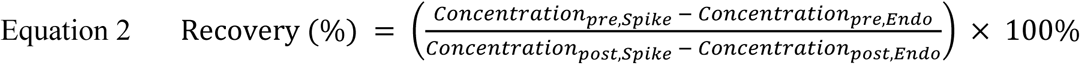

### Donor Analysis

Fecal samples from individual donors (n = 10) were prepared in triplicate according to the procedure for preparing quality-control samples (vide supra), with the modification that 30–50 mg of lyophilized fecal material was used per sample. Additionally, 10-fold dilutions were prepared by diluting 30 µL of each sample with 270 µL of 20% aqueous acetonitrile to account for analytes that may be present in amounts above the upper limit of quantitation (ULOQ). Samples were subjected to LC–MS/MS analysis immediately after preparation.

### LC–MS/MS Instrumentation and Parameters

Sample aliquots of 150 µL were added to a PlateOne round-bottom 96-well plate, sealed with NAL-96 Sealing Film (USA Scientific, Ocala, FL), and loaded into a Waters Sample Organizer refrigerated to 10 °C. A 2 µL aliquot of each sample was injected via a flow-through needle and Acquity I-Class Premier UPLC. The analytes were separated with a Waters Acquity Premier BEH C_18_ column (50 mm by 2.1 mm, 1.7 µm particle size), equipped with a VanGuard FIT BEH C_18_ guard column, and maintained at 50 °C. Solvent A was DDI H_2_O + 0.1% acetic acid, and Solvent B was acetonitrile + 0.1% acetic acid with a flow-rate of 0.5 mL/min. To provide an effective balance between signal intensity, peak shape, peak separation, and total run-time, the solvent gradient was 20% B to 40% B from 0–3 min, then ramped to 98% B from 3–4 min, held isocratic at 98% B from 4–4.5 min, reduced to 2% B from 4.5–4.9 min, and equilibrated to 20% B from 4.9–5.0 min.

Targeted mass spectrometry was performed with a Xevo TQ Absolute triple quadrupole mass spectrometer (Waters, Milford, MA, USA), operated in multiple-reaction monitoring (MRM) and negative electrospray ionization (ESI^—^) modes. Spectra were collected using MassLynx 4.2. The sample eluted through the capillary of the ZSpray source set to -0.80 kV and was sprayed toward the source cone set to 150 °C, with N_2_ desolvation gas flowing at 800 L/hr and 600 °C. The precursor ion, fragment ion, cone voltage, and collision energy (**Table 1**) were determined by injecting each analyte individually at 10 µM in acetonitrile, with each parameter selected to maximize the signal intensity for each analyte. Data were processed with Waters’ software module, TargetLynx.

**Table 1:** Mass Spectrometric Parameters. Mass spectrometric parameters for the selective quantitation of all analytes and internal standards using multi-reaction monitoring in negative-ion mode. Q1 and Q3 are the precursor and fragment ions’ m/z, respectively. CV is the cone voltage, and CE is the collision energy.

| **Phenolic Acids** | Q1<br>(m/z) | Q3<br>(m/z) | CV<br>(V) | CE<br>(eV) |
| --- | --- | --- | --- | --- |
| 3-(3-Hydroxyphenyl)<br>propanoic Acid (3,3-HPPA) | 165.0 | 121.0 | 20 | 20 |
| 3-(4-Hydroxyphenyl)<br>propanoic Acid (3,4-HPPA) | 165.0 | 93.0 | 20 | 10 |
| Caffeic Acid (CFA) | 180.0 | 135.0 | 10 | 20 |
| <sup>13</sup>C<sub>3</sub>-Caffeic Acid | 181.9 | 136.9 | 10 | 20 |
| Dihydrocaffeic Acid (DCFA) | 182.0 | 137.0 | 20 | 10 |
| Dihydroferulic Acid (DFA) | 195.0 | 136.0 | 20 | 20 |
| Ferulic Acid (FA) | 193.0 | 134.0 | 10 | 20 |
| *para*-Coumaric Acid (pCA) | 163.0 | 119.0 | 10 | 20 |
| Protocatechuic Acid (CAT) | 152.9 | 108.9 | 20 | 20 |
| Sinapinic Acid (SIN) | 223.0 | 92.9 | 20 | 30 |
| Vanillic Acid (VANAC) | 167.0 | 107.9 | 10 | 20 |
| **Lignans** |  |  |  |  |
| Hydroxymatairesinol (HMAT) | 373.0 | 173.0 | 10 | 30 |
| Lariciresinol (LARI) | 359.0 | 329.0 | 10 | 10 |
| Matairesinol (MAT) | 357.0 | 83.0 | 30 | 30 |
| Pinoresinol (PINO) | 357.0 | 121.0 | 20 | 20 |
| Secoisolariciresinol (SECO) | 361.0 | 165.0 | 20 | 20 |
| d<sub>6</sub>-Secoisolariciresinol | 367.2 | 167.9 | 20 | 30 |
| **Enterolignans** |  |  |  |  |
| Enterodiol (END) | 301.0 | 253.0 | 10 | 25 |
| Enterolactone (ENL) | 297.0 | 107.0 | 20 | 30 |
| <sup>13</sup>C<sub>3</sub>-Enterolactone | 300.0 | 254.9 | 20 | 20 |

## RESULTS AND DISCUSSION

### Method Validation

#### Sensitivity

The chromatographic conditions were optimized to yield the lowest possible LLOQ for each analyte. The phenolic acids were quantified with LLOQ’s between 0.24 nM and 3.91 nM, and the lignans and enterolignans were quantified with LLOQ’s between 0.24 nM and 0.98 nM (**Table 2**). These results meet or exceed similar LC–MS/MS methods, where many phenolic acids have been reported with LLOQ’s between 19.5 nM^34^ and 800.4 nM,^45^ and many lignans have been reported with LLOQ’s between 0.068 nM^46^ and 3.91 nM.^31^ Carryover was found to be negligible for all analytes, ranging from 0.01%–0.07%.

**Table 2:** Method Sensitivity. Summary of key sensitivity metrics for each compound. L is the lower limit of quantitation, U is the upper limit of quantitation, C is the percent carryover, and R^2^ is the coefficient of determination from linear regression.

| **Phenolic Acids** | **L**<br>**(nM)** | **U**<br>**(nM)** | **C**<br>**(%)** | **R<sup>2</sup>** |
| --- | --- | --- | --- | --- |
| 3-(3-Hydroxyphenyl)<br>propanoic Acid (3,3-HPPA) | 1.95 | 500 | 0.037 | 0.998 |
| 3-(4-Hydroxyphenyl)<br>propanoic Acid (3,4-HPPA) | 0.98 | 500 | 0.017 | 0.997 |
| Caffeic Acid (CFA) | 3.91 | 500 | 0.066 | 0.999 |
| Dihydrocaffeic Acid (DCFA) | 3.91 | 500 | 0.003 | 0.998 |
| Dihydroferulic Acid (DFA) | 3.91 | 500 | 0.042 | 0.997 |
| Ferulic Acid (FA) | 0.25 | 500 | 0.008 | 0.996 |
| *para*-Coumaric Acid (pCA) | 0.98 | 500 | 0.027 | 0.998 |
| Protocatechuic Acid (CAT) | 1.95 | 500 | 0.070 | 0.998 |
| Sinapinic Acid (SIN) | 0.98 | 500 | 0.020 | 0.998 |
| Vanillic Acid (VANAC) | 0.25 | 500 | 0.029 | 0.998 |
| **Lignans** |  |  |  |  |
| Hydroxymatairesinol (HMAT) | 0.49 | 500 | 0.010 | 0.997 |
| Lariciresinol (LARI) | 0.98 | 500 | 0.023 | 0.996 |
| Matairesinol (MAT) | 0.25 | 500 | 0.011 | 0.998 |
| Pinoresinol (PINO) | 0.98 | 500 | 0.040 | 0.999 |
| Secoisolariciresinol (SECO) | 0.49 | 500 | 0.016 | 0.997 |
| **Enterolignans** |  |  |  |  |
| Enterodiol (END) | 0.25 | 500 | 0.006 | 0.998 |
| Enterolactone (ENL) | 0.25 | 500 | 0.015 | 0.997 |

Various solvent conditions were tested to achieve the best balance between peak shape and signal intensity. Neat solvents produced the largest signal intensities for all analytes, but they led to split-peaks for most phenolic acids (**Figure 2**). Adding 0.1% formic acid to the chromatographic solvents caused a loss of signal intensity, such that the LLOQ for each analyte was 5–100 times higher than with either neat solvents or with 0.1% acetic acid. This is exemplified by the signal intensity for dihydroferulic acid (DHFA), which was highest in neat solvents across the entire calibration range (**Figure 2A**), but the linearity was poor due to peak splitting (**Figure 2B**). Similarly, the signal for pinoresinol (PINO) was obliterated when 0.1% formic acid was used in the chromatographic solvents, but the use of 0.1% acetic acid allowed for a strong, unambiguous peak at 0.31 nM (**Figure 2C–D**). These results recapitulate the finding of Song et al. that 0.1% acetic acid added to chromatographic solvents, instead of additives like formic acid, ammonium acetate, or ammonium fluoride, yields the best ion efficiencies and fragment ion coverage of various phenolic acids and lignin oligomers.^47^

**Figure 2:**
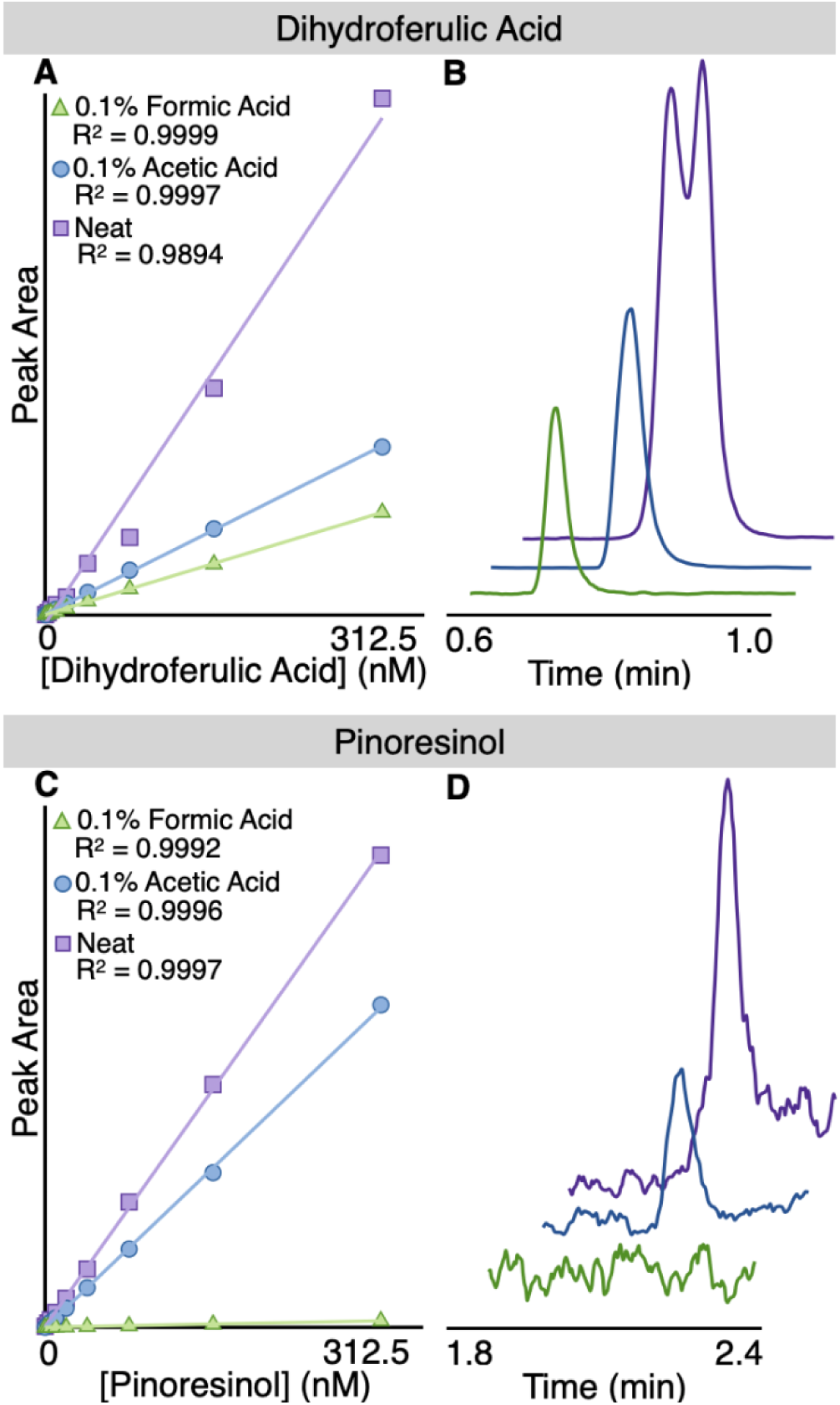
Chromatographic Solvent Additives Impact Peak Shape and Signal Intensity. The mobile phase mixture of water and acetonitrile was tested alone (neat) or with additives (0.1% formic acid or 0.1% acetic acid). The representative effects of these additives on peak detection is depicted for **(A–B)** dihydroferulic acid and **(C–D)** pinoresinol. **(A,C)** Calibration curves and **(B,D)** extracted-ion chromatograms (dihydroferulic acid, 312.5 nM; pinoresinol, 0.31 nM) are shown.

#### Selectivity

The chromatographic conditions were optimized to selectively identify and quantify each analyte. While there was overlap in the elution of some phenolic acids, their distinct precursor and fragment ions allowed for unique identification (**Figure S2A**). There was no overlap in the peaks for the lignans and enterolignans (**Figure S2B**). The analytes 3-(3-hydroxyphenyl)propanoic acid (3,3-HPPA) and pinoresinol (PINO) display cross-signal contributions from other analytes, but baseline separation allows for accurate identification and quantification. The peak at 0.64 minutes that appears in the extracted-ion chromatogram of 3,3-HPPA results from a fragment of 3-(4-hydroxyphenyl)propanoic acid (3,4-HPPA) because they have the same precursor m/z. To confirm this, each compound was injected separately. When 3,4-HPPA was injected alone, a signal appeared in the channel that is configured to detect 3,3-HPPA (**Figure S2C**). Similarly, pinoresinol showed a second peak at 2.3 minutes in its extracted-ion chromatogram that is the result of a cross-signal contribution from matairesinol (**Figure S2D**).

#### Matrix Effects

The matrix effects in this method are generally negligible, with most analytes displaying no more than 5% ion suppression or enhancement (**Table S1**). These results correspond to the matrix factor calculated by Nørskov et al. in their detection of lignans from porcine fecal samples^40^ and by Sánchez-Patán et al. in their quantification of phenolic acids from fecal waters.^34^ Ion enhancements of 76.0% and 51.1% were observed for 3,3-HPPA at the 10 nM and 40 nM concentrations, respectively; one of the replicates had a high amount of 3,3-HPPA endogenous to the fecal sample (286 nM), which challenged accurate quantification of the samples spiked with 10 nM and 40 nM 3,3-HPPA. Similarly, 3,4-HPPA was impacted by ion enhancements of 38.2% and 37.4% at 10 nM and 40 nM, respectively. Nonetheless, the matrix effects for these analytes at the 100 nM and 400 nM concentrations were no more than 8.2%. Ion suppression of 16.5% was observed for protocatechuic acid at the 10 nM concentration, whereas González-Domínguez observed a 24% ion enhancement when protocatechuic acid was spiked into a plasma matrix that was extracted with acetonitrile containing 1.5 M formic acid and 10 mM ammonium formate.^31^

Background contaminants imparted by polypropylene microcentrifuge tubes (MCT’s) are a concern in LC–MS/MS analyses because they can lead to inaccurate quantification, whether by ion suppression, ion enhancement, or by cross-signal contribution. Multiple studies have found that MCT’s can impart small molecule contaminants that are used as antistatic agents,^48^ plastic stabilizers,^49^ and plastic clarifying agents into biological samples.^50^ Canez and Li found that using MCT’s for lipidomics imparted more contaminants than did using borosilicate glass.^51^ We have developed this method around the use of glass during sample preparation because the use of MCT’s during liquid-liquid extraction gave rise to unpredictable, strong signals with the same precursor m/z, fragment m/z, and retention time as some analytes in the panel (**Figure 3** and **Figure S3**).

**Figure 3:**
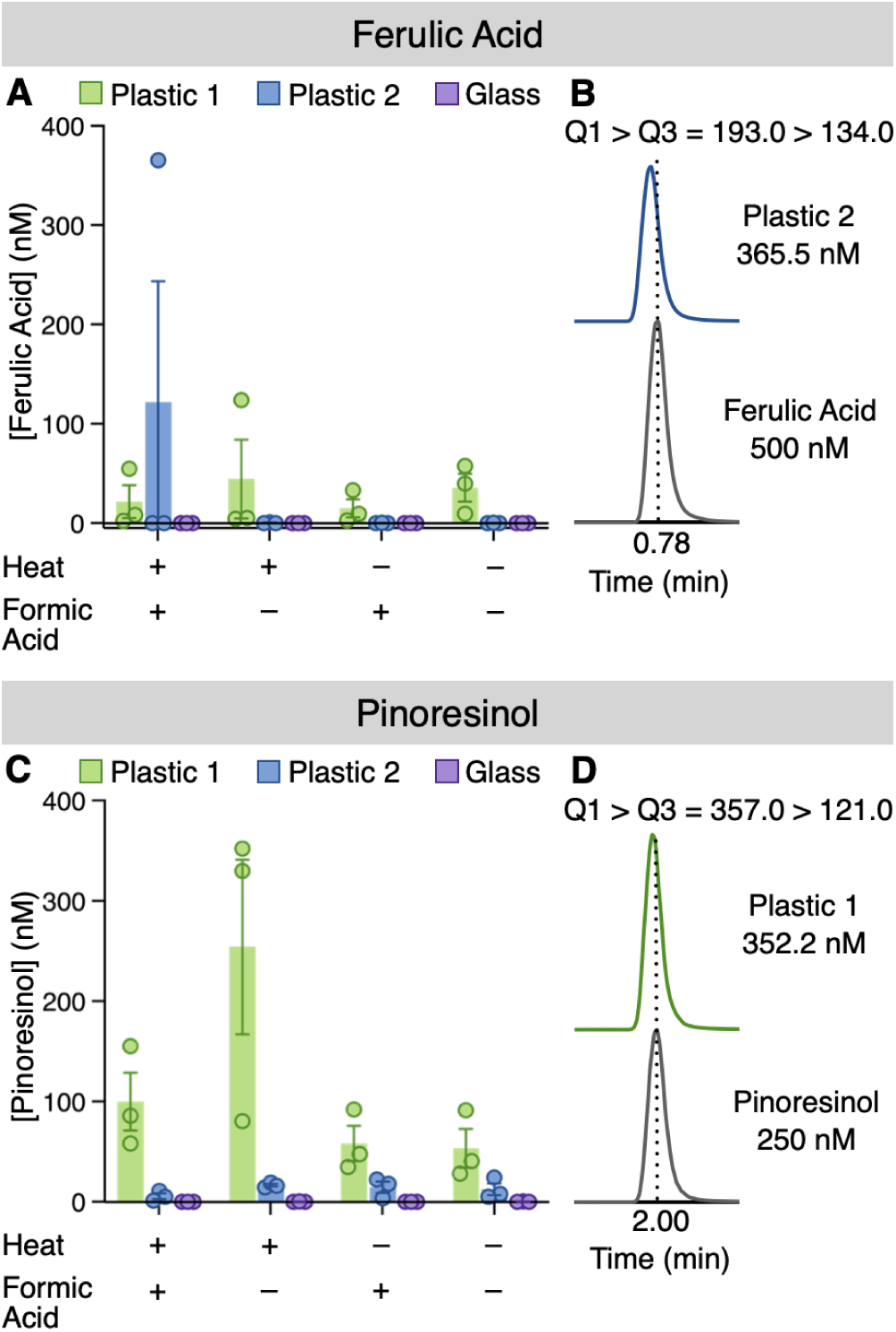
Extraction Vessel Impacts Background Signal. Leachate from two different brands of polypropylene microcentrifuge tubes (MCT’s) upon liquid-liquid extraction of water, with or without heat and 0.1% formic acid, using methyl *tert*-butyl ether. For **(A–B)** ferulic acid and **(C–D)** pinoresinol comparison of the observed concentrations of **(A)** ferulic acid or **(C)** pinoresinol after extraction from either plastic or glass. Extracted-ion chromatograms of MCT extracts are shown in comparison to standards of **(B)** ferulic acid or **(D)** pinoresinol. n = 3 biological replicates; bars denote means ± S.E.M.

We tested an array of sample processing conditions with MCT’s from two different manufacturers against glass vials, each in triplicate. MCT leachate from Brand 1 produced signals in the ferulic acid channel between 2.5 nM and 123.8 nM, across all processing conditions. Brand 2 MCT leachate produced a signal in the ferulic acid channel equivalent to 365.5 nM when DDI H_2_O was heated in MCT’s and extracted with MtBE containing 0.1% formic acid (**Figure 3A**). The chromatograms in **Figure 3B** compare the absolute ferulic acid signal from a 500 nM standard to the signal observed when DDI H_2_O in MCT’s from Brand 2 was heated and extracted with MtBE containing 0.1% formic acid. Leachate from MCT’s also impacted pinoresinol. Brand 1 MCT’s produced a signal of 352.2 nM in the pinoresinol channel when DDI H_2_O was heated and extracted using MtBE without formic acid. Brand 1 produced signals that ranged from 28.0 nM to 92.1 nM across the remaining 3 treatment conditions: DDI H_2_O heated and extracted using MtBE + 0.1% formic acid, DDI H_2_O extracted using MtBE + 0.1% formic acid, and DDI H_2_O extracted using MtBE. Leachate from Brand 2 MCT’s produced signals in the pinoresinol channel that ranged from 1.1 nM to 24.1 nM across all conditions (**Figure 3C–D**). In MCT’s, adding water and extracting with MtBE caused interfering signals for other phenolic acids and enterolignans that ranged from 2–10 times their respective LLOQ (**Figure S3**). This phenomenon has not previously been reported for the analysis of phenolic acids, lignans, or enterolignans. The use of glass vials and culture tubes during sample preparation alleviated this problem.

#### Accuracy, Precision, and Stability

Accuracy, precision, and stability are crucial validation parameters because they help to establish that a method is robust and replicable. All analytes were quantified with accuracy and precision results within the acceptable tolerance (± 20% at the LLOQ and ± 15% for all other concentrations), per FDA guidelines,^41^ except for vanillic acid (**Table 3**). The results for most of the 17 analytes were within ± 10% relative error and relative standard deviation, which matches the results of similar methods,^34,40^ with a few exceptions. Relative standard deviations for precision were generally high, yet acceptable, for protocatechuic acid (10.2% to 18.9%), due to the large endogenous amount of this analyte in fecal samples. The accuracy of vanillic acid was acceptable at the lowest concentration tested (-9.2% relative error at 10 nM), but it was between -20.2% and -26.4% relative error at 40 nM, 100 nM, and 400 nM. Sánchez-Patán reported a relative error of 13.33% for the accuracy of vanillic acid but a -20.00% relative error for caffeic acid accuracy using stable-isotope labelled 4-hydroxybenzoic acid as an internal standard.^34^ Conversely, the method reported herein substantially improved accuracy for caffeic acid (relative error range: -1.1%–2.7%) by using stable-isotope labelled caffeic acid as an internal standard. Together, these data highlight that the choice of IS can impact the accuracy and precision of a method. The accuracy and precision that this method affords was recapitulated in stability validation experiments; allowing samples to rest on the benchtop for up to 72 hours or to be subjected to freeze-thaw before analysis did not impact the accuracy and precision of this method (**Table S2**).

**Table 3:** Robust Accuracy, Precision, and Recovery Metrics in Human Fecal Sample Matrix. Human fecal samples were spiked with standards at A = 10 nM, B = 40 nM, C = 100 nM, or D = 400 nM. For accuracy and precision measurements, n = 6 independent sample sets, with n = 6 replicate injections per sample set. Accuracy is reported as relative error (RE) and precision is reported as relative standard deviation (RSD). For recovery measurements, n = 3 independent sample sets, with n = 3 replicate injections per sample set.

| | Accuracy<br>(RE%) | | | | Precision<br>(RSD%) | | | | Recovery (%)<br>Mean ± S.E.M | | | |
| --- | --- | --- | --- | --- | --- | --- | --- | --- | --- | --- | --- | --- |
|  | A | B | C | D | A | B | C | D | A | B | C | D |
| **Phenolic Acids** |  |  |  |  |  |  |  |  |  |  |  |  |
| 3-(3-Hydroxyphenyl)propanoic Acid | 2.9 | 5.1 | 1.2 | -2.8 | 7.7 | 5.8 | 7.5 | 5.3 | 101.7 ± 3.1 | 105.5 ± 0.6 | 105.2 ± 0.6 | 102.5 ± 0.6 |
| 3-(4-Hydroxyphenyl)propanoic Acid | 0.4 | 4.6 | 2.8 | -2.1 | 8.6 | 5.3 | 7.9 | 5.1 | 94.2 ± 2.7 | 100.2 ± 1.2 | 102.8 ± 0.8 | 102.4 ± 1.0 |
| Caffeic Acid | 1.4 | 2.7 | -0.3 | -1.1 | 11.1 | 5.8 | 7.9 | 6.4 | 93.9 ± 1.0 | 98.9 ± 0.4 | 99.7 ± 0.4 | 96.1 ± 0.5 |
| Dihydrocaffeic Acid | -6.0 | -2.8 | -5.7 | -9.3 | 11.3 | 8.8 | 9.2 | 7.5 | 79.6 ± 2.9 | 91.3 ± 1.7 | 92.3 ± 1.2 | 94.3 ± 1.3 |
| Dihydroferulic Acid | -3.1 | -3.6 | -6.6 | -11.4 | 9.9 | 8.1 | 8.2 | 6.8 | 99.9 ± 2.0 | 100.5 ± 0.4 | 101.0 ± 0.6 | 98.4 ± 0.6 |
| Ferulic Acid | 3.9 | 4.7 | 0.9 | -2.9 | 11.6 | 6.9 | 8.8 | 6.7 | 82.9 ± 4.4 | 100.0 ± 1.1 | 101.4 ± 0.7 | 100.9 ± 0.5 |
| *para*-Coumaric Acid | 1.4 | 4.9 | 2.7 | 0.5 | 6.0 | 4.0 | 7.9 | 6.4 | 100.8 ± 0.8 | 103.3 ± 0.6 | 104.1 ± 0.4 | 104.0 ± 0.6 |
| Protocatechuic Acid | 0.0 | 6.2 | 6.1 | 5.1 | 18.9 | 10.2 | 10.6 | 11.0 | 100.4 ± 15.8 | 106.8 ± 5.1 | 106.3 ± 3.0 | 107.9 ± 3.3 |
| Sinapinic Acid | -3.4 | 0.1 | -2.4 | -7.2 | 10.8 | 7.9 | 9.4 | 8.0 | 89.9 ± 1.5 | 90.7 ± 0.4 | 92.0 ± 1.1 | 90.2 ± 0.8 |
| Vanillic Acid | -9.2 | -20.2 | -24.3 | -26.4 | 12.7 | 5.0 | 6.7 | 4.4 | 81.2 ± 6.8 | 81.3 ± 9.7 | 91.0 ± 3.7 | 102.5 ± 2.2 |
| **Lignans** |  |  |  |  |  |  |  |  |  |  |  |  |
| Hydroxymatairesinol | 4.9 | 9.7 | 7.6 | 5.2 | 6.9 | 4.1 | 5.6 | 4.7 | 106.0 ± 1.7 | 98.9 ± 2.0 | 98.4 ± 2.2 | 98.2 ± 1.1 |
| Lariciresinol | 12.3 | 11.5 | 8.8 | 7.6 | 5.7 | 4.7 | 4.7 | 4.4 | 109.2 ± 1.8 | 104.2 ± 2.1 | 105.3 ± 2.3 | 103.8 ± 1.2 |
| Matairesinol | 5.3 | -0.8 | 0.1 | -3.6 | 10.0 | 6.3 | 9.0 | 5.1 | 102.1 ± 1.2 | 98.2 ± 0.8 | 100.3 ± 0.8 | 98.8 ± 0.9 |
| Pinoresinol | 4.3 | -1.2 | -0.6 | -3.4 | 7.7 | 6.3 | 7.3 | 6.0 | 92.0 ± 4.4 | 98.3 ± 1.0 | 95.9 ± 1.8 | 99.3 ± 0.9 |
| Secoisolariciresinol | 0.5 | 1.2 | -1.1 | -1.4 | 8.6 | 4.9 | 5.8 | 2.1 | 101.2 ± 1.8 | 98.4 ± 2.3 | 97.3 ± 1.9 | 96.9 ± 0.9 |
| **Enterolignans** |  |  |  |  |  |  |  |  |  |  |  |  |
| Enterodiol | 5.3 | -3.1 | -4.1 | -3.8 | 8.1 | 6.5 | 5.0 | 3.6 | 101.4 ± 1.3 | 97.5 ± 1.5 | 98.6 ± 0.9 | 98.1 ± 1.3 |
| Enterolactone | 6.0 | 2.1 | 0.2 | -0.9 | 6.8 | 2.4 | 5.4 | 2.9 | 97.4 ± 2.3 | 99.4 ± 0.9 | 100.2 ± 0.6 | 100.4 ± 0.7 |

#### Recovery

Recovery is an important validation parameter because it reflects the efficacy of the sample processing steps: if the recovery is too low, then improvements are required to ensure that the entire amount of each analyte is being measured, and if the recovery is too high, then there may be an inflationary impact from matrix-associated ion-enhancement phenomena.^52^ The recoveries for this method generally range from 95% to 105% (**Table 3**). The recovery of sinapinic acid was low, averaging 90.7% across all spiked concentrations, while the recoveries for protocatechuic acid and lariciresinol were slightly high, between 105% and 110%. The recovery of vanillic acid in samples spiked with 10 nM and 40 nM of this analyte was 81.2% and 81.3%, respectively—an unsurprising finding due to the relatively low accuracy with which this analyte was measured.

### Human Fecal Sample Analysis

Diets that are high in polyphenols, including phenolic acids and lignans, are associated with reduced risks for cancer^36^ and cardiovascular disease,^53^ which motivated the development of this assay. Thus, we tested the applicability of our method to the analysis of human fecal samples from 10 healthy donors (**Table 4**). The analyte that reached the highest concentration across the fecal samples tested was the monophenol 3,3-HPPA (range: 4.82–98.60 nmol/g of lyophilized fecal material; median: 19.3 ± 26.8 nmol/g of lyophilized fecal material). Indeed, the amounts of 3,3-HPPA as well as 3,4-HPPA in the samples from one donor were greater than the upper limit of quantitation for this method. The measured gut microbiome abundance of 3,3-HPPA is corroborated by Gutiérrez-Díaz et al. who reported that, out of 30 phenolic acids, 3,3-HPPA had the fourth highest concentration in fecal samples from a cohort of Spanish adults who ate a Mediterranean diet.^54^ Gonthier et al. found that 3,3-HPPA was increased 15-fold in human urine following consumption of a polyphenol-rich diet,^33^ and Lu et al. found that 3,3-HPPA was a metabolic end product when human fecal inoculum was fermented with ferulic acid.^55^ Collectively, our finding that 3,3-HPPA is an abundant phenolic acid in human fecal samples aligns with others’ quantifications of this analyte, supporting the applicative utility of our method.

**Table 4:** Analytes are Effectively Measured in Human Fecal Samples. Measured concentrations of phenolic acids, lignans, and enterolignans in human fecal samples (n = 10 independent donors). Values are mean ± S.E.M. (nmol/g), normalized to the dry mass of each sample; n = 3 technical replicates per donor. ULOQ denotes the upper limit of quantitation, and “nd” denotes not detected.

| Phenolic acids | Human Fecal Sample Donor |  |  |  |  |  |  |  |  |  |
| --- | --- | --- | --- | --- | --- | --- | --- | --- | --- | --- |
|  | 1 | 2 | 3 | 4 | 5 | 6 | 7 | 8 | 9 | 10 |
| 3-(3-Hydroxyphenyl)propanoic Acid | 4.82 ± 0.21 | 21.99 ± 0.14 | 11.75 ± 0.21 | 98.60 ± 0.91 | 14.83 ± 0.57 | 39.10 ± 2.18 | 9.89 ± 0.05 | >ULOQ | 28.16 ± 1.97 | 15.13 ± 0.13 |
| 3-(4-Hydroxyphenyl)propanoic Acid | 1.82 ± 0.18 | 6.35 ± 0.10 | 2.97 ± 0.06 | 15.39 ± 0.08 | 5.36 ± 0.10 | 11.94 ± 0.42 | 3.69 ± 0.07 | >ULOQ | 7.42 ± 0.09 | 2.59 ± 0.03 |
| Caffeic Acid | 7.32 ± 0.28 | 8.80 ± 0.22 | 9.09 ± 0.62 | 8.76 ± 0.36 | 8.94 ± 0.46 | 11.89 ± 0.63 | 13.95 ± 2.00 | 8.83 ± 0.63 | 13.26 ± 0.35 | 7.87 ± 0.13 |
| Dihydrocaffeic Acid | 4.09 ± 0.08 | 9.06 ± 0.14 | 3.98 ± 0.15 | 7.65 ± 0.23 | 5.01 ± 0.15 | 12.11 ± 1.15 | 3.63 ± 0.08 | 11.57 ± 0.27 | 9.03 ± 0.21 | 8.94 ± 0.11 |
| Dihydroferulic Acid | 10.06 ± 0.39 | 8.07 ± 0.39 | 2.54 ± 0.09 | 2.79 ± 0.07 | 9.08 ± 0.18 | 11.95 ± 3.82 | 5.18 ± 0.13 | 4.28 ± 0.17 | 2.53 ± 0.05 | 1.61 ± 0.03 |
| Ferulic Acid | 12.47 ± 0.59 | 9.62 ± 0.63 | 4.59 ± 0.38 | 5.06 ± 0.34 | 33.48 ± 0.52 | 11.44 ± 0.57 | 7.93 ± 0.39 | 9.09 ± 0.55 | 3.89 ± 0.21 | 2.36 ± 0.09 |
| *para*-Coumaric Acid | 6.13 ± 0.44 | 3.31 ± 0.06 | 3.50 ± 0.24 | 2.23 ± 0.08 | 3.45 ± 0.05 | 9.72 ± 0.15 | 2.78 ± 0.13 | 17.57 ± 1.95 | 3.18 ± 0.07 | 1.44 ± 0.07 |
| Protocatechuic Acid | 15.65 ± 0.99 | 39.53 ± 0.58 | 8.75 ± 0.34 | 19.31 ± 0.67 | 27.65 ± 5.14 | 17.92 ± 0.67 | 15.34 ± 0.61 | 11.60 ± 0.29 | 40.78 ± 1.82 | 19.67 ± 0.42 |
| Sinapinic Acid | 2.11 ± 0.16 | 2.21 ± 0.02 | 1.28 ± 0.09 | 0.72 ± 0.02 | 3.22 ± 0.04 | 4.59 ± 0.13 | 1.23 ± 0.02 | 0.76 ± 0.05 | 1.21 ± 0.02 | 0.44 ± 0.01 |
| Vanillic Acid | 2.79 ± 0.21 | 5.36 ± 0.16 | 4.31 ± 0.17 | 8.45 ± 0.17 | 2.15 ± 0.05 | 11.03 ± 0.44 | 6.23 ± 0.26 | 3.13 ± 0.07 | 6.72 ± 0.14 | 1.42 ± 0.03 |
| **Lignans** |  |  |  |  |  |  |  |  |  |  |
| Hydroxymatairesinol | 0.10 ± 0.00 | 0.09 ± 0.00 | 0.22 ± 0.03 | 0.69 ± 0.05 | 0.10 ± 0.00 | 0.42 ± 0.01 | 0.10 ± 0.02 | 0.16 ± 0.00 | 0.08 ± 0.00 | 0.23 ± 0.02 |
| Lariciresinol | 0.20 ± 0.01 | 0.20 ± 0.02 | 0.89 ± 0.06 | 0.55 ± 0.04 | 0.21 ± 0.01 | 0.93 ± 0.05 | 0.45 ± 0.08 | 0.62 ± 0.03 | 0.56 ± 0.02 | 1.31 ± 0.03 |
| Matairesinol | 0.17 ± 0.00 | 0.16 ± 0.00 | 0.08 ± 0.00 | 0.61 ± 0.02 | 0.38 ± 0.01 | 0.78 ± 0.02 | 0.08 ± 0.00 | 0.65 ± 0.01 | 0.10 ± 0.01 | 0.05 ± 0.00 |
| Pinoresinol | 0.20 ± 0.00 | 1.70 ± 0.23 | 0.73 ± 0.06 | 0.16 ± 0.01 | 0.21 ± 0.04 | 3.23 ± 0.63 | 0.49 ± 0.04 | 0.49 ± 0.05 | 0.56 ± 0.09 | 0.17 ± 0.00 |
| Secoisolariciresinol | nd | 0.12 ± 0.01 | 0.11 ± 0.01 | 0.29 ± 0.02 | 0.21 ± 0.02 | 0.43 ± 0.02 | 2.18 ± 0.30 | 0.89 ± 0.11 | 0.21 ± 0.00 | 0.28 ± 0.01 |
| **Enterolignans** |  |  |  |  |  |  |  |  |  |  |
| Enterodiol | 4.15 ± 0.13 | 5.21 ± 0.16 | 1.81 ± 0.07 | 34.51 ± 1.84 | 2.38 ± 0.06 | 8.80 ± 0.43 | 7.32 ± 1.19 | 11.08 ± 1.24 | 4.35 ± 0.10 | 1.39 ± 0.03 |
| Enterolactone | 11.45 ± 0.75 | 6.36 ± 0.35 | 9.97 ± 0.19 | 123.67 ± 2.83 | 3.33 ± 0.22 | 35.56 ± 2.69 | 23.54 ± 0.24 | 110.45 ± 1.53 | 15.62 ± 0.24 | 111.12 ± 3.34 |

The phenolic acids with the second and third highest median ± interquartile range of concentrations across fecal samples were the catechols protocatechuic acid (range: 8.75–40.78 nmol/g of lyophilized fecal material; median: 18.2 ± 7.7 nmol/g of lyophilized fecal material), and caffeic acid (range: 7.32–13.95 nmol/g of lyophilized fecal material; median: 9.1 ± 3.5 nmol/g of lyophilized fecal material). In addition to catechols being prevalent in dietary plants, these molecules can also be synthesized in the gut microbiome from corresponding guaiacol precursors via demethylation. Such guaiacol/catechol pairs composing our analyte panel include ferulic acid/caffeic acid, dihydroferulic acid/dihydrocaffeic acid, and vanillic acid/protocatechuic acid. For these analytes, we summed the guaiacols and catechols and found that 6 of 10 donors (Donors 2, 3, 4, 8, 9, and 10) harbored significantly greater amounts of catechols than guaiacols (**Figure 4**). Conversion of guaiacols to catechols entails demethylation of a 3-methoxy, 4-hydroxyphenyl moiety to reveal 3,4-dihydroxyphenyl with enhanced redox capacity. Specifically, the vicinal hydroxyl groups can form a chelate with iron and, thereby, undergo oxidation. The ability of catechols to effectively chelate iron has implicated this class of molecules in disease processes that depend on iron and its oxidation state.^56^ Thus, the abundance of catechols in the gut relative to guaiacols may be a valuable predictor of a gut microbiome’s oxidation potential and corresponding ability to modulate disease processes.

**Figure 4:**
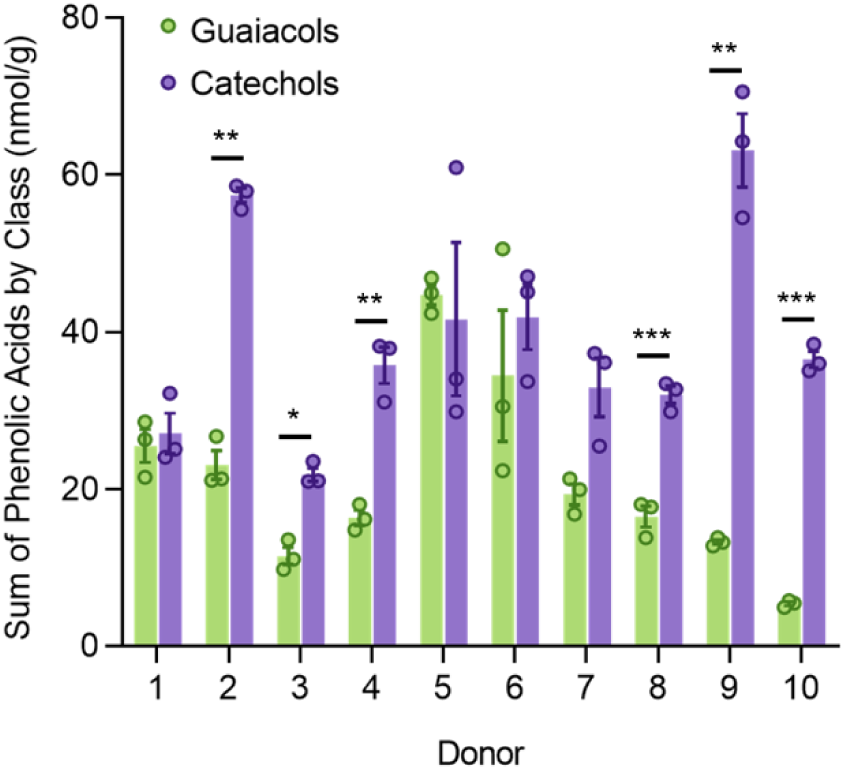
Differences in Total Phenolic Acid Content, by Class. Sum of guaiacols (dihydroferulic acid, ferulic acid, vanillic acid) and catechols (caffeic acid, dihydrocaffeic acid, protocatechuic acid) measured in lyophilized fecal samples each from 10 different human donors. n = 3 technical replicates; bars denote means ± S.E.M.; significance was determined by paired t-test; ***: P < 0.001, **: P < 0.01, *: P < 0.05.

Quantification of lignans and enterolignans revealed lignans to be in relatively low abundance (0.05–3.23 nmol/g of lyophilized fecal material), whereas the enterolignans were up to 38-fold more abundant (enterodiol, 1.39–34.51 nmol/g of lyophilized fecal material; enterolactone, 3.33–123.67 nmol/g of lyophilized fecal material) (**Table 4**). The increased abundance of enterolignans in comparison with lignans aligns with known biosynthetic pathways. Specifically, the 5 lignans in our panel are biochemical precursors to the enterolignans via processing by human gut bacteria.^27^ Kasimir et al. incubated bacteria from pig ceca with uniformly ^13^C-labelled lignin and found that lignin-derived secoisolariciresinol was converted to enterolactone within 8 hours and depleted within 24 hours.^57^ While many existing methods either do not report concentrations for some lignans or they report that some lignans were not detected,^38,40^ the method reported here enabled quantification of all 5 lignans and 2 enterolignans in samples from all 10 donors.

In conclusion, the validated LC–MS/MS method described here has been designed to be simple and robust, with the goal of facilitating gut microbiome research that seeks to elucidate the roles of polyphenolic metabolites in human health and disease. This method advances the field of phenolic acid, lignan, and enterolignan quantitation in human fecal samples by simultaneously quantifying these diet- and gut microbiome-derived metabolites with a short analytical cycle of 5 minutes per injection. Importantly, this method allows for direct comparisons between individuals and across time because the analyte concentrations are normalized to the mass of lyophilized fecal material, thereby removing any confounding effects imparted by differences in water content. Taken together, this accurate, sensitive method to quantify molecules that are linked to disease prevention can enable future work that tailors their gut bacterial biosynthesis for the benefit of human health.

## ASSOCIATED CONTENT

## Supporting Information

Photos of mixing apparatus to facilitate processing of samples in 2 mL glass HPLC vials (Figure S1); extracted-ion chromatograms for each analyte and extracted-ion chromatograms for 3-(3-hydroxyphenyl)propanoic acid and pinoresinol showing cross-signal contributions (Figure S2); summary of results for validation experiments to investigate matrix effects (Table S1); interference of molecules leached from plastic microcentrifuge tubes on analyte signals (Figure S3); summary of results for validation experiments to investigate stability (Table S2).

## AUTHOR INFORMATION

## Corresponding Author

Elizabeth N. Bess **–** Department of Chemistry, University of California, Irvine, California 92617, United States; Department of Molecular Biology and Biochemistry, University of California, Irvine, California 92617, United States;

## Author Contributions

CAD and ENB designed experiments. CAD, DNC, JDR, and ENB conducted data analysis. CAD, DNC, and JDR performed experiments. CAD and ENB wrote and edited the manuscript. ENB acquired funds and provided project supervision and administration. All authors read and approved the final version of the manuscript.

## Note

The authors declare no competing financial interests.

## ACKNOWLEDGMENTS

This study was made possible in part through access to the Mass Spectrometry Core Facility of the Department of Chemistry, a shared resource at the University of California, Irvine. We appreciated the contributions of facility director Dr. Felix Grün who provided technical assistance and experimental guidance. The table of contents graphic was created with BioRender.com (agreement number *CV28OZ0DCB*).

