## Supplemental Figures for "A Validated LC–MS/MS Method for Quantifying Phenolic Acids, Lignans, and Enterolignans from Human Fecal Samples"

### SUPPORTING INFORMATION

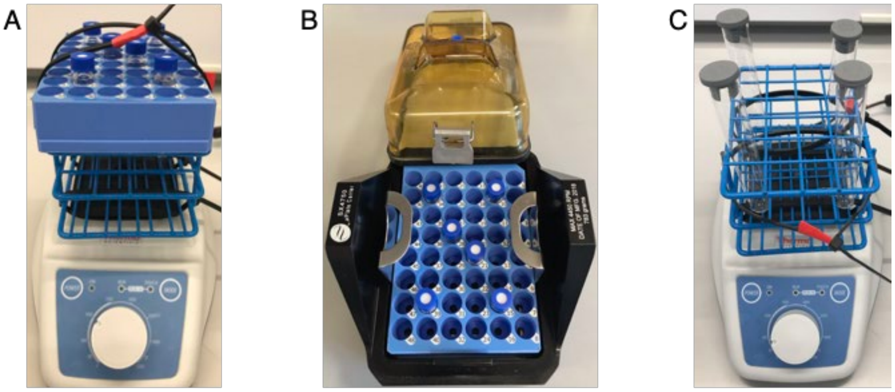


**Figure S1: Apparatus for Mixing and Centrifugation of Samples in 2 mL Glass HPLC Vials or Glass Culture Tubes. (A)** A 24-slot tube rack was affixed to a Thermo Scientific LP Vortex Mixer with cable ties. For mixing of samples in HPLC vials, a 48-well HPLC autosampler vial rack was placed on top of the tube rack and held in place with additional cable ties. **(B)** For centrifugation, the 48-well HPLC autosampler vial rack was inserted into the microplate carriers for a Beckman-Coulter Avanti J-15R benchtop centrifuge. **(C)** Glass culture tubes capped with butyl rubber stoppers were mixed at 500 rpm in a 24-slot tube rack affixed to a Thermo Scientific LP Vortex Mixer.

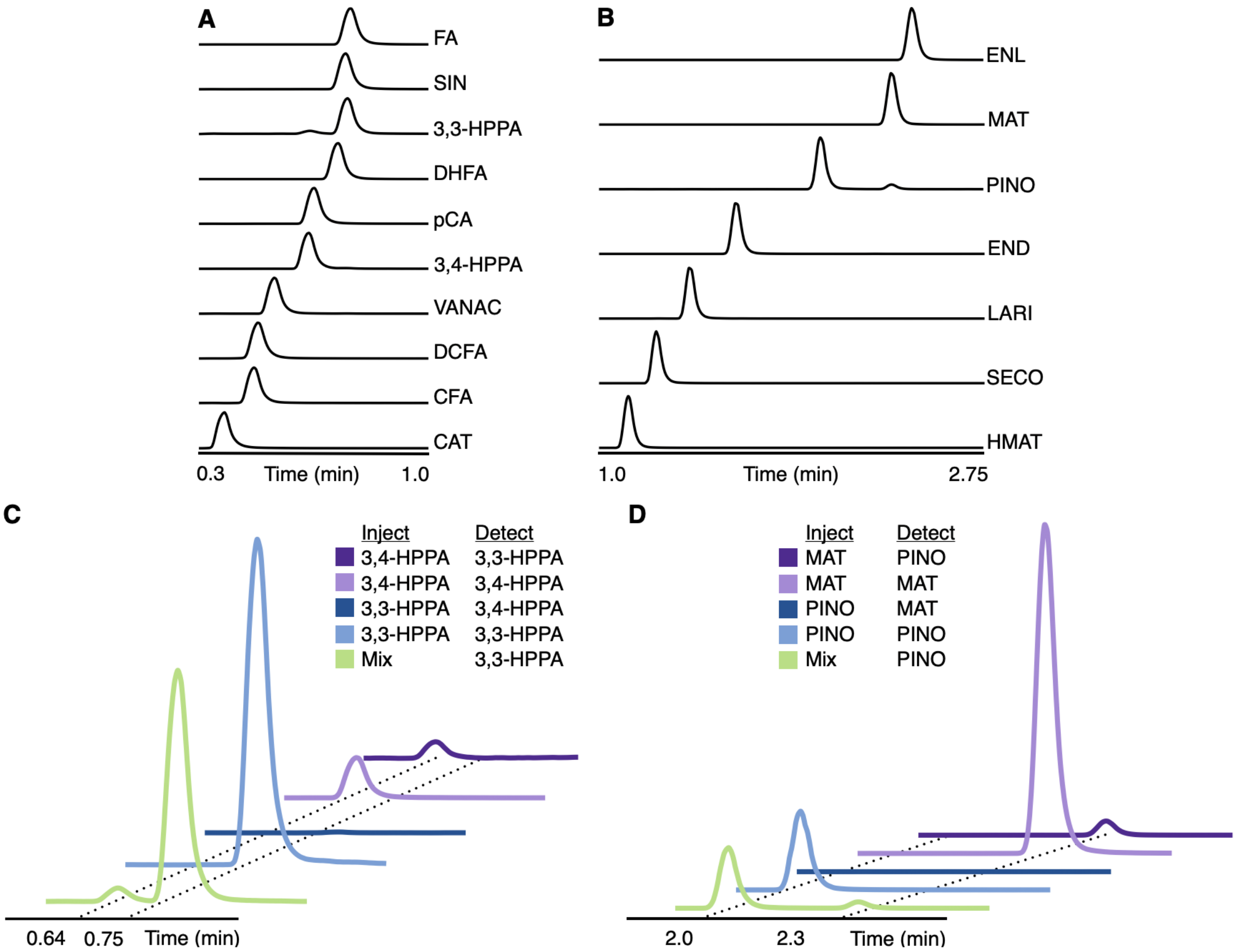


**Figure S2: Cross-Signal Contributions Across Analytes.** Extracted-ion chromatograms of all 17 analytes quantified with this method and extracted-ion chromatograms showing cross-signal contributions. **(A)** Overlaid extracted-ion chromatograms of all phenolic acids, monitored from 0.3–1.0 minutes. **(B)** Overlaid extracted-ion chromatograms of 3-(3-hydroxyphenyl)propanoic acid (3,3-HPPA) and 3-(4-hydroxyphenyl)propanoic acid (3,4-HPPA), showing that the peak at 0.64 minutes in the extracted-ion chromatogram for 3,3-HPPA from a mixed sample of 3,3-HPPA and 3,4-HPPA is a cross-signal contribution from 3,4-HPPA. **(C)** Overlaid extracted-ion chromatograms for all lignans and enterolignans. **(D)** Overlaid extracted-ion chromatograms of pinoresinol (PINO) and matairesinol (MAT), showing that the peak at 2.3 minutes in the extracted-ion chromatogram for pinoresinol from a mixed sample of pinoresinol and matairesinol is a cross-signal contribution from matairesinol. "Detect" means that the mass spectrometer was set to the parameters for detection of the analyte listed in the "Detect" column.

**Table S1: Matrix Effects.** Matrix effects impacting analytes in human fecal samples spiked with standards at A = 10 nM, B = 40 nM, C = 100 nM, or D = 400 nM compared to mixtures of analytes in neat solutions at the same nominal concentrations. Results are the percent difference of the spiked analyte response to the neat analyte response. Values are mean ± S.E.M.; n = 3 independent sample sets, with n = 3 replicate injections.

| **Phenolic Acids** | A (%) | B (%) | C (%) | D (%) |
|---|---|---|---|---|
| 3-(3-Hydroxyphenyl)propanoic Acid | 76.0 ± 40.6 | 51.1 ± 19.7 | 8.2 ± 4.7 | 3.1 ± 3.5 |
| 3-(4-Hydroxyphenyl)propanoic Acid | 38.2 ± 24.2 | 37.4 ± 15.8 | 7.1 ± 3.8 | 2.1 ± 3.4 |
| Caffeic Acid | 0.8 ± 4.9 | -4.8 ± 1.2 | 0.4 ± 1.1 | -1.6 ± 0.6 |
| Dihydrocaffeic Acid | -11.3 ± 5.7 | 2.8 ± 1.4 | 1.1 ± 1.2 | 4.1 ± 1.1 |
| Dihydroferulic Acid | -6.1 ± 2.4 | 0.9 ± 1.3 | -2.3 ± 0.9 | 1.8 ± 1.4 |
| Ferulic Acid | -3.4 ± 1.1 | 0.8 ± 1.6 | 2.8 ± 0.7 | 3.6 ± 1.4 |
| *para*-Coumaric Acid | -6.5 ± 0.5 | -4.8 ± 0.8 | -4.4 ± 0.9 | -2.8 ± 2.0 |
| Protocatechuic Acid | -16.5 ± 2.8 | 4.8 ± 2.2 | -1.2 ± 0.8 | -0.8 ± 2.5 |
| Sinapinic Acid | -2.2 ± 1.0 | 3.3 ± 1.1 | 1.9 ± 0.9 | 7.1 ± 1.4 |
| Vanillic Acid | -16.9 ± 0.8 | -6.2 ± 1.0 | -6.3 ± 0.9 | -1.9 ± 2.4 |
| **Lignans** | | | | |
| Hydroxymatairesinol | -2.6 ± 1.0 | -3.3 ± 0.9 | 0.2 ± 0.6 | -0.5 ± 0.8 |
| Lariciresinol | 1.2 ± 1.2 | 0.6 ± 1.1 | 4.9 ± 1.4 | 2.1 ± 1.1 |
| Matairesinol | -1.9 ± 0.6 | 1.3 ± 0.9 | 0.1 ± 0.6 | -0.1 ± 0.6 |
| Pinoresinol | 0.0 ± 0.0 | -0.8 ± 0.8 | -0.3 ± 1.2 | 1.1 ± 1.1 |
| Secoisolariciresinol | 3.0 ± 1.1 | 0.8 ± 0.9 | 4.5 ± 1.0 | 1.7 ± 0.6 |
| **Enterolignans** | | | | |
| Enterodiol | 4.0 ± 0.9 | 5.3 ± 0.8 | 3.5 ± 0.7 | 4.8 ± 0.7 |
| Enterolactone | 0.4 ± 2.1 | 2.8 ± 1.0 | -0.8 ± 0.7 | 0.1 ± 0.4 |

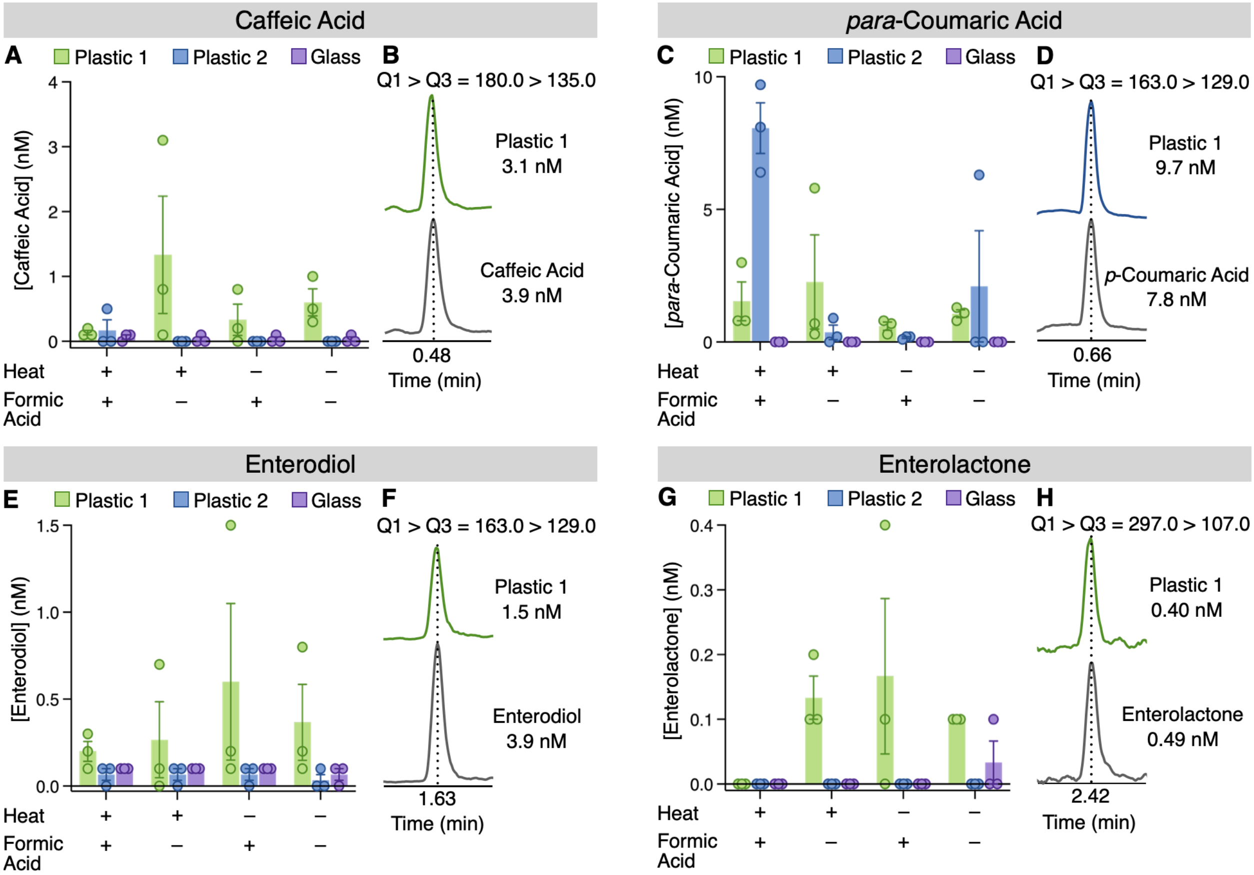


**Figure S3: Extraction Vessel Impacts Background Signal.** Leachate from two different brands of polypropylene microcentrifuge tubes (MCT's) upon liquid-liquid extraction of water, with or without heat and 0.1% formic acid, using methyl *tert*-butyl ether. For **(A–B)** caffeic acid, **(C–D)** *para*-coumaric acid, **(E–F)** enterodiol, and **(G–H)** enterolactone, comparison of the observed concentrations of **(A)** caffeic acid, **(C)** *para*-coumaric acid, **(E)** enterodiol, and **(G)** enterolactone after extraction from either plastic or glass. Extracted-ion chromatograms of MCT extracts are shown in comparison to standards of **(B)** caffeic acid, **(D)** *para*-coumaric acid, **(F)** enterodiol, or **(H)** enterolactone. n = 3 biological replicates; bars denote means ± S.E.M.

**Table S2: Benchtop and Freeze-Thaw Stability.** Accuracy and precision of analytes in human fecal samples spiked with standards at A = 10 nM, B = 40 nM, C = 100 nM, or D = 400 nM, and either left to sit on the benchtop or stored at -20 °C for between 12 hr and 72 hr. Accuracy is reported as relative error (RE) and precision is reported as relative standard deviation (RSD); n = 3 independent sample sets, with n = 6 replicate injections per sample set.

| | Benchtop Stability | | | | | | | | Freeze-Thaw Stability | | | | | | | |
|---|---|---|---|---|---|---|---|---|---|---|---|---|---|---|---|---|
| | Accuracy (RE%) | | | | Precision (RSD%) | | | | Accuracy (RE%) | | | | Precision (RSD%) | | | |
| **Phenolic Acids** | A | B | C | D | A | B | C | D | A | B | C | D | A | B | C | D |
| 3-(3-Hydroxyphenyl) propanoic Acid | 4.9 | 7.7 | 6.3 | -0.3 | 4.6 | 2.9 | 5.1 | 3.4 | 4.4 | 6.5 | 0.7 | -2.9 | 4.8 | 3.5 | 6.0 | 2.3 |
| 3-(4-Hydroxyphenyl) propanoic Acid | 8.4 | 4.5 | 7.0 | 0.4 | 6.3 | 5.9 | 4.6 | 3.6 | 0.7 | 4.5 | -0.8 | -1.2 | 7.4 | 6.9 | 7.3 | 4.6 |
| Caffeic Acid | -3.0 | -1.5 | -4.6 | -7.1 | 14.2 | 5.7 | 4.6 | 3.3 | 1.5 | 2.4 | -4.8 | -3.7 | 3.6 | 2.8 | 4.9 | 1.6 |
| Dihydrocaffeic Acid | -3.4 | -3.2 | -4.8 | -9.9 | 6.8 | 5.0 | 4.0 | 3.9 | 3.9 | 0.6 | -4.5 | -9.6 | 7.4 | 7.7 | 6.9 | 2.4 |
| Dihydroferulic Acid | 4.0 | 2.8 | 2.9 | -6.8 | 9.5 | 5.8 | 7.8 | 5.0 | -4.9 | -0.4 | -9.8 | -10.2 | 4.3 | 5.1 | 2.8 | 4.1 |
| Ferulic Acid | 7.4 | 5.0 | 4.5 | -1.9 | 9.5 | 2.7 | 4.1 | 1.2 | 3.2 | 8.5 | -0.8 | -2.4 | 5.6 | 3.8 | 6.8 | 1.8 |
| *para*-Coumaric Acid | 3.5 | 4.0 | 4.0 | -0.3 | 6.6 | 3.7 | 3.5 | 4.4 | 1.8 | 7.2 | 2.2 | 0.2 | 4.5 | 4.2 | 5.3 | 1.6 |
| Protocatechuic Acid | -15.2 | -1.1 | 4.1 | -1.9 | 28.6 | 8.2 | 6.7 | 4.8 | 1.8 | 15.4 | -0.9 | -0.5 | 10.6 | 38.7 | 9.4 | 5.7 |
| Sinapinic Acid | 6.9 | 0.1 | -1.5 | -6.1 | 6.6 | 5.1 | 4.7 | 2.3 | -1.8 | -2.9 | -8.2 | -7.8 | 4.0 | 2.4 | 4.3 | 2.5 |
| Vanillic Acid | -2.0 | -13.7 | -16.0 | -19.9 | 13.1 | 4.2 | 4.5 | 6.3 | -11.8 | -16.7 | -26.4 | -28.6 | 21.0 | 15.9 | 12.8 | 14.1 |
| **Lignans** | | | | | | | | | | | | | | | | |
| Hydroxymatairesinol | -0.4 | -0.4 | -3.0 | -4.7 | 10.5 | 8.8 | 11.0 | 7.0 | -10.7 | -6.8 | -8.4 | -7.6 | 4.5 | 5.3 | 5.1 | 2.1 |
| Lariciresinol | 11.1 | 5.7 | 4.9 | 2.4 | 4.1 | 4.0 | 7.3 | 8.7 | 7.9 | 4.7 | 2.2 | 3.9 | 5.9 | 6.2 | 8.5 | 4.6 |
| Matairesinol | 11.2 | 5.9 | 4.5 | -0.2 | 4.1 | 6.2 | 8.1 | 5.5 | 8.6 | 2.8 | -1.7 | -2.1 | 3.4 | 6.6 | 3.6 | 5.1 |
| Pinoresinol | 11.3 | 7.3 | 8.4 | 5.8 | 6.6 | 5.4 | 6.9 | 6.5 | 9.8 | -4.0 | -1.6 | -1.6 | 10.1 | 6.6 | 11.1 | 9.0 |
| Secoisolariciresinol | 2.5 | 0.3 | -0.1 | -1.7 | 4.1 | 4.9 | 5.9 | 3.9 | -1.1 | -1.2 | -3.4 | -0.2 | 3.5 | 6.7 | 6.4 | 2.6 |
| **Enterolignans** | | | | | | | | | | | | | | | | |
| Enterodiol | 6.9 | 3.6 | 2.0 | -0.1 | 8.0 | 2.7 | 2.8 | 3.8 | 5.8 | 0.2 | -6.2 | -3.3 | 9.7 | 6.2 | 3.3 | 6.6 |
| Enterolactone | 3.5 | 2.2 | 1.4 | -3.0 | 5.1 | 4.5 | 5.3 | 2.8 | 4.2 | -0.1 | -5.1 | -11.5 | 6.7 | 3.0 | 5.5 | 1.8 |
